# Temporal transitions of spontaneous brain activity

**DOI:** 10.1101/166512

**Authors:** Zhiwei Ma, Nanyin Zhang

**Author notes:** **Address for correspondence:** Dr. Nanyin Zhang, Hartz Family Professor, Department of Biomedical Engineering and Electrical Engineering, The Huck Institutes of Life Sciences, The Pennsylvania State University, W-341 Millennium Science Complex, University Park, PA 16802, USA. Conflict of interest: none.

## Abstract

Spontaneous brain activity, typically investigated using resting-state fMRI (rsfMRI), provides a measure of inter-areal resting-state functional connectivity (RSFC). Previous rsfMRI studies mainly focused on spatial characteristics of RSFC, but the temporal relationship between RSFC patterns is still elusive. Particularly, it remains unknown whether separate RSFC patterns temporally fluctuate in a random manner, or transit in specific orders. Here we investigated temporal transitions between characteristic RSFC patterns in awake rats and humans. We found that transitions between RSFC patterns were reproducible and significantly above chance, suggesting that RSFC pattern transitions were nonrandom. The organization of RSFC pattern transitions in rats was analyzed using graph theory. Pivotal RSFC patterns in transitions were identified including hippocampal, thalamic and striatal networks. This study has revealed nonrandom temporal relationship between characteristic RSFC patterns in both rats and humans. It offers new insights into understanding the spatiotemporal dynamics of spontaneous activity in the mammalian brain.

## Introduction

Multiple lines of evidence indicate that spontaneous brain activity plays an essential role in brain function (Raichle ME and MA Mintun 2006; Zhang D and ME Raichle 2010). For instance, intrinsic neuronal signaling consumes the vast majority of brain energy (Raichle ME 2006, 2010). Investigation of spontaneous brain activity, predominantly conducted by resting-state functional magnetic resonance imaging (rsfMRI) (Biswal B et al. 1995; Fox MD and ME Raichle 2007), has provided critical insight into the intrinsic organization of the brain network. Using spontaneously fluctuating blood-oxygenation-level dependent (BOLD) signal measured by rsfMRI, resting-state functional connectivity (RSFC) between brain regions can be gauged by statistical interdependence of their rsfMRI signals over the period of data acquisition (Fox MD and ME Raichle 2007). Based on this quantity, multiple brain networks of functionally-related regions have been identified in both humans and animals, which convey the information of stable functional connectivity organization of the brain (Beckmann CF et al. 2005; Fox MD et al. 2005; Damoiseaux JS et al. 2006; Smith SM et al. 2009; Allen EA et al. 2011; Liang Z et al. 2011).

Conventional rsfMRI studies generally focus on steady features of RSFC by assuming that RSFC is stationary during the acquisition period. However, meaningful temporal variability of RSFC at shorter time scales within rsfMRI scans has also been discovered (Chang C and GH Glover 2010). This initial research and its follow-up studies revealed dynamic properties of RSFC, indicating that the stationarity assumption of RSFC would be overly simplistic for understanding spontaneous brain activity (Hutchison RM, T Womelsdorf, EA Allen, et al. 2013; Preti MG et al. 2016). Indeed, using sliding window analysis and clustering methods, temporally alternating but spatially repeatable RSFC patterns have been identified (Allen EA et al. 2014). In addition, Liu and Duyn (2013) developed a method that examined instantaneous co-activations of BOLD signal at single rsfMRI frames and found that BOLD co-activation patterns well corresponded brain connectivity configurations (Liu X and JH Duyn 2013). With this method, the default mode network, which is a single network under the assumption of stationary RSFC, can be decomposed into multiple sub-networks with distinct spatiotemporal characteristics and functional relevance (Liu X and JH Duyn 2013). Notably, the neurophysiologic relevance of dynamic RSFC has been validated in multiple studies using simultaneous electrophysiology and rsfMRI acquisitions (Tagliazucchi E et al. 2012; Chang C et al. 2013; Keilholz SD 2014).

In parallel with blossoming dynamic RSFC studies in humans, dynamic RSFC studies in animal models have also been conducted. Animals’ brain preserves fundamental organizational properties as the human brain (Liang Z *et al.* 2011; Ma Z et al. 2016), and can serve as a translational model for studying complicated brain dynamics. Using either the sliding window or co-activation pattern approach, dynamic RSFC patterns have been found in both awake and anesthetized rats, as well as in anesthetized monkeys (Majeed W et al. 2011; Hutchison RM, T Womelsdorf, JS Gati, et al. 2013; Keilholz SD et al. 2013; Mohajerani MH et al. 2013; Liang Z et al. 2015; Grandjean J et al. 2017; Ma Y et al. 2017). These results suggest that dynamics in RSFC might be a general feature in the mammalian brain.

Despite the critical advancement, aforementioned dynamic rsfMRI studies have mainly focused on revealing the *spatial* patterns of dynamic RSFC, while the *temporal relationship* between these RSFC patterns is still unclear. Particularly, it remains elusive whether characteristic brain connectivity patterns temporally fluctuate in a random manner, or evolve in specific sequences (Majeed W *et al.* 2011; Zalesky A et al. 2014; Mitra A et al. 2015; Preti MG *et al.* 2016). Lack of such information highlights a gap in elucidating the temporal organization of separate brain connectivity configurations, and thus hinders comprehensive characterization of spatiotemporal dynamics of spontaneous brain activity.

To address this issue, in the present study we studied the temporal transitions of intrinsic brain activity in awake rats and humans. We used a library of characteristic RSFC patterns as the references (Gordon EM et al. 2016; Ma Z *et al.* 2016). Each rsfMRI frame was matched to one of the reference RSFC patterns that had the highest spatial similarity to the BOLD co-activation pattern of the frame (Liu X and JH Duyn 2013; Liang Z *et al.* 2015). This step generated a time series of characteristic RSFC patterns for each rsfMRI run. Temporal transitions between every pair of RSFC patterns were then counted, which created a RSFC pattern transition network. The reproducibility of the RSFC pattern transition networks in rats and humans was separately examined. In addition, a weighted direct graph constructed based on the RSFC pattern transition network in rats was constructed, and its topological organization was studied using graph theory analysis. RSFC patterns that were pivotal in temporal transitions were further identified.

## Materials and methods

### Animals

41 Long-Evans (LE) adult male rats were used. Data from 31 rats were also used in another study (Ma Z *et al.* 2016) and were reanalyzed for the purpose of the present study. All rats were housed in Plexiglas cages with controlled ambient temperature (22-24 °C) and maintained on a 12 h light:12 h dark schedule. Food and water were provided ad libitum. The experiment was approved by the Institutional Animal Care and Use Committee (IACUC) at the Pennsylvania State University.

### Rat MRI experiments

Rats were acclimated to the MRI environment for seven days following the procedures described in (Zhang N et al. 2010; Liang Z *et al.* 2011, 2012, 2012, 2014; Gao YR et al. 2016) to minimize motion and stress. For the setup of awake animal imaging, the rat was first briefly anesthetized with 2-3% isoflurane and fit into a head holder with a built-in coil and a body tube. Isoflurane was discontinued and the rat was placed into the magnet. All rats were fully awake during imaging. A similar approach has also been used for awake rodent fMRI in other groups (Bergmann E et al. 2016; Chang PC et al. 2016; Yoshida K et al. 2016).

MRI data acquisition was conducted on a Bruker 7T small animal MRI scanner (Billerica, MA). Anatomical MRI images were acquired using a T1-weighted rapid imaging with refocused echoes (RARE) sequence with the following parameters: repetition time (TR) = 1500 ms; echo time (TE) = 8 ms; matrix size = 256 × 256; field of view (FOV) = 3.2 × 3.2 cm^2^; slice number = 20; slice thickness = 1 mm; RARE factor = 8. rsfMRI images were acquired using a T2*-weighted gradient-echo echo planar imaging (EPI) sequence with the following parameters: TR = 1000 ms; TE =15 ms; matrix size = 64 × 64; FOV = 3.2 × 3.2 cm^2^; slice number = 20; slice thickness= 1 mm. 600 EPI volumes were acquired for each run, and two to four runs were acquired for each animal.

### Rat image preprocessing

Detailed description of the image preprocessing pipeline can be found in (Ma Z *et al.* 2016) and is briefly summarized as follows. Relative framewise displacement (FD) (Power JD et al. 2012) of rat brain EPI images were calculated, and EPI volumes with FD > 0.2 mm and their immediate temporal neighbors were removed (1.75% of total rsfMRI volumes). The first 10 volumes of each rsfMRI run were also removed to warrant a steady state of magnetization. Brain normalization to a standard rat brain was performed using Medical Image Visualization and Analysis (MIVA, <u>http://ccni.wpi.edu/). Head motion was corrected using SPM12 (http://www.fil.ion.ucl.ac.uk/spm/</u>). In-plane spatial smoothing was carried out using a Gaussian filter (FWHM = 0.75 mm). Nuisance regression was performed with the regressors of three translation and three rotation motion parameters estimated by SPM as well as white matter and ventricle signals. Band-pass filtering was performed with the frequency band of 0.01–0.1 Hz.

### Characteristic RSFC patterns

To obtain a library of characteristic RSFC spatial patterns in the awake rat brain, we used a RSFC-based whole-brain parcellation scheme (40 parcels) we previously published (Ma Z *et al.* 2016). In this scheme, voxels with similar RSFC patterns were grouped together, so that RSFC patterns were similar within parcels but dissimilar across parcels (Ma Z *et al.* 2016). As a result, these 40 RSFC patterns represented a set of characteristic RSFC patterns in the awake rat brain and were used as the references (also see Supplemental Information).

All characteristic RSFC patterns were obtained using seed-based correlational analysis with each parcel as the seed. Specifically, the regionally-averaged time course from all voxels within the seed region was used as the seed time course, and the Pearson cross-correlation coefficient between the seed time course and the time course of each individual brain voxel was calculated. Correlation analysis was performed for the first 540 volumes of each rsfMRI run to ensure the same degree of freedom. Correlation coefficients were then Fisher's Z-transformed. For each parcel, its group-level RSFC map was voxelwise calculated using one-sample t-test based on a linear mixed-effect model with the random effect of rats and the fixed effect of Z values for each run. The spatial similarity between these reference RSFC patterns was determined by pairwise spatial correlations between every two characteristic RSFC patterns.

### Temporal transitions between RSFC patterns

To analyze temporal transitions between RSFC patterns, a time sequence of framewise RSFC patterns (1 sec each frame) was first obtained by matching each rsfMRI frame to one of the 40 reference RSFC patterns, based on the notion that BOLD co-activation patterns in single rsfMRI frames also represent their RSFC patterns (Liu X et al. 2013; Liu X and JH Duyn 2013; Liang Z *et al.* 2015). To do so, preprocessed rsfMRI time series were first demeaned and variance normalized. Subsequently, the spatial Pearson correlation coefficients between each rsfMRI frame and individual reference RSFC patterns in the library were respectively calculated. The reference RSFC pattern that best matched the rsfMRI frame (i.e. the reference RSFC pattern that had the highest spatial correlation) was selected. To ensure the correspondence between each rsfMRI frame and the matched RSFC pattern was statistically meaningful, we set a minimal threshold of the spatial correlation coefficient > 0.1 (p value < 10^−13^). 89.9% of total volumes met this criterion. Frames that did not meet this criterion (10.09% of total volumes) were labeled as not corresponding to any reference RSFC patterns. This step generated a time sequence of framewise RSFC patterns. In this sequence, each rsfMRI frame was denoted by a number between 1 and 40, representing its correspondence to one of the 40 reference RSFC patterns. The number 0 was used to denote rsfMRI frames not corresponding to any reference RSFC patterns, as well as frames removed in image preprocessing (e.g. frames with large FD). In the sequence, the number of transitions between every two RSFC patterns was counted (i −> j, where i ≠ j, i ≠ 0 and j ≠ 0). Transitions involving 0 (i.e., 0 −> 0, or 0 −> i, or i −> 0, where i ≠ 0) were not counted. This procedure yielded a 40 × 40 RSFC pattern transition matrix, where its entry (i, j) represented the number of transitions between RSFC pattern i to pattern j.

### Reproducibility of temporal transitions between RSFC patterns

The reproducibility of temporal transitions between RSFC patterns was assessed at both the group and individual levels. At the group level, we used a split-group approach. All 41 rats were randomly divided into two subgroups with 20 rats in subgroup 1 and 21 rats in subgroup 2. The RSFC pattern transition matrix was computed for each subgroup. Entries in each matrix were normalized to the range of [0, 1], and the correlation of the corresponding off-diagonal matrix entries between the two subgroups was assessed.

It is possible that spatially similar RSFC patterns have a higher chance to transit between each other in both subgroups, and this systematic bias might inflate the reproducibility of RSFC pattern transitions between the two subgroups. To control for this effect, we regressed out the spatial similarities between every two reference RSFC patterns, quantified by their spatial correlation values, from the transition matrices in both subgroups and then assessed the reproducibility again.

Reproducibility of temporal transitions between RSFC patterns was also evaluated at the individual level. For each rat, its individual-level transition matrix was obtained, and the reproducibility was computed using Pearson correlation of the corresponding off-diagonal matrix entries between this individual-level transition matrix and the group-level transition matrix.

### Organization of RSFC pattern transitions

The group-level transition matrix was thresholded to identify transitions that were statistically significant. The p value of each entry in the transition matrix was calculated using the permutation test. Since we are only interested in transitions between two different RSFC patterns, before the permutation test, the temporal sequence of RSFC patterns was consolidated by combining consecutively repeated appearances of the same pattern to one appearance of the pattern. For example, four consecutive appearances of Pattern ‘x’ (i.e. ‘xxxx’) were replaced by one ‘x’. This consolidated temporal sequence was then permuted 10000 times, and a transition matrix was obtained for each permuted sequence. This step generated an empirical null distribution for each off-diagonal entry in the transition matrix, and the p value of the entry was obtained accordingly. p values were further adjusted using false-discovery rate (FDR) correction at the FDR rate of 0.05 (Genovese CR et al. 2002). Entries with insignificant p values were set to zero. All entries were then rescaled to the range of [0,1]. Finally, similarities between RSFC patterns were regressed out from nonzero entries.

Using this thresholded transition matrix as the adjacency matrix, a graph was constructed using Gephi 0.9.1 (https://gephi.org/). In this weighted directed graph, each node represented a RSFC pattern, and each edge connecting two nodes signified an above-chance transition between two RSFC patterns with the edge weight proportional to the normalized number of transitions.

Graph theory analysis of this RSFC pattern transition network was performed using the Brain Connectivity Toolbox (https://sites.google.com/site/bctnet/). The community affiliation of nodes in the graph was obtained by repeating the Louvain community detection algorithm (Vincent DB et al. 2008) for 1000 times to ensure a stable solution. Specifically, for each repetition, a 40 x 40 matrix was generated so that its entry (i, j) was 1 if nodes i and j were in the same community and 0 otherwise. The average of these 1000 matrices was then binarized using a threshold of 0.9, and the final inference of the community affiliation was obtained from the node affiliation of connected components in the binarized matrix (Liang Z *et al.* 2011).

To identify the hub nodes in the transition graph, local graph measures of node strength, betweenness centrality, local characteristic path length and local clustering coefficient of each node were first computed. Using these node metrics, hub nodes with high node strength, high betweenness centrality, short distance to other nodes, and low local clustering coefficient (Bullmore E and O Sporns 2009) were identified using the method described in (van den Heuvel MP et al. 2010). Briefly, a hub score (0 to 4) was given to each node according to the total number of the following criteria the node met: (1) upper 20 percentile in node strength; (2) upper 20 percentile in betweenness centrality; (3) lower 20 percentile in characteristic path length; and (4) lower 20 percentile in local clustering coefficient. Node met at least three criteria was defined as a hub, indicating its pivotal role in transitions between RSFC patterns.

### Reproducibility of RSFC pattern temporal transitions in the human brain

The reproducibility of temporal transitions between RSFC patterns in the human brain was evaluated using a similar process. The human data used were the ‘extensively preprocessed 3T rsfMRI data’ from 812 subjects, which were a subset of the S1200 Subjects Data Release of the Human Connectome Project (HCP, https://www.humanconnectome.org/) (Van Essen DC et al. 2013). All rsfMRI data were acquired on a 3T Siemens Skyra MRI scanner using a multi-band EPI sequence with the parameters of TR = 720 ms, TE = 33.1 ms, flip angle = 52°, FOV = 208 × 180 mm^2^, matrix size = 104 × 90, voxel size = 2 × 2 × 2 mm^3^, slice number = 72, slice thickness = 2 mm, multiband factor = 8 (Feinberg DA et al. 2010; Moeller S et al. 2010; Setsompop K et al. 2012; Glasser MF et al. 2013). Data preprocessing used the HCP minimal preprocessing pipelines (Glasser MF *et al.* 2013), MSM-All brain registration (Robinson EC et al. 2014), and the ICA+FIX pipeline (Griffanti L et al. 2014; Salimi-Khorshidi G et al. 2014), and these procedures were completed by the HCP.

To obtain a library of characteristic RSFC patterns in the human brain, we used a well-established RSFC-based parcellation scheme (333 parcels) (Gordon EM *et al.* 2016), which has been demonstrated to have high within-parcel homogeneity and reflect the underlying connectivity structure of the human brain (Gordon EM *et al.* 2016). Based on this scheme, 333 characteristic RSFC patterns were obtained using seed-based correlational analysis with individual parcels as seeds. For each rsfMRI run, the seed time course was averaged from all grayordinates within the seed, and Pearson cross-correlation coefficient between the seed time course and the time course of each individual cortical grayordinate was calculated. Correlation coefficients were Fisher's Z-transformed. For each parcel, its group-level RSFC map was grayordinate-wise calculated by one-sample t-test using a linear mixed-effect model with the random effect of subjects and the fixed effect of Z values for individual runs. Pairwise spatial correlations between these group-level RSFC maps were also calculated to measure their similarity.

Each rsfMRI frame was matched to one of the 333 reference patterns that had the highest spatial similarity to the BOLD co-activation pattern of the frame, gauged by their spatial Pearson correlation. The minimal spatial correlation coefficient was set at 0.05. rsfMRI frames below this threshold were labeled as not corresponding to any reference RSFC patterns. This step generated a temporal sequence of RSFC patterns for each rsfMRI run. In this sequence, each rsfMRI frame was denoted by a number between 1 and 333, representing its correspondence to one of the 333 characteristic RSFC patterns. The number 0 was used to denote rsfMRI frames not corresponding to any reference RSFC patterns. In the sequence, the number of transitions between every two RSFC patterns was counted (i −> j, where i ≠ j, i ≠ 0 and j ≠ 0). Transitions involving 0 (i.e., 0 −> 0, or 0 −> i, or i −> 0, where i ≠ 0) were not counted. This procedure yielded a 333 × 333 temporal transition matrix for each run. The temporal transition matrix for each subject was obtained by summing the temporal transition matrices from all 4 rsfMRI runs. The group-level transition matrix was obtained by averaging the subject-level temporal transition matrices across all subjects.

The reproducibility of temporal transitions between RSFC patterns in the human brain was also assessed at both the individual and group levels. The individual-level reproducibility was calculated based on the correlation of off-diagonal entries between the individual’s transition matrix and the group-level transition matrix. The group-level reproducibility was evaluated using a split-group approach. All 812 subjects were randomly divided into two subgroups (406 subjects each subgroup). The RSFC pattern transition matrix was computed for each subgroup, respectively, and the correlation of off-diagonal matrix entries between the two subgroups were assessed. To control for the effect of the spatial similarity between characteristic RSFC patterns, we also regressed out the spatial correlation values between characteristic RSFC patterns from the transition matrices in both subgroups and then re-assessed the reproducibility.

## Results

In this study, we investigated the temporal transitions between spontaneous brain activity patterns in awake rats and humans. In rat data, we first obtained a library of characteristic RSFC patterns, using seed-based correlational analysis with seeds defined by parcels in a whole-brain RSFC-based parcellation. These characteristic RSFC patterns were used as the reference patterns. Subsequently, based on the notion that the BOLD co-activation patterns of single rsfMRI frames represent their RSFC patterns (Liu X *et al.* 2013; Liu X and JH Duyn 2013), each rsfMRI frame was matched to one of the 40 reference RSFC patterns based on its BOLD co-activation pattern. This step generated a time sequence of characteristic RSFC patterns for each rsfMRI run. Temporal transitions between every pair of RSFC patterns were then counted, which provided a RSFC pattern transition matrix. A weighted directed transition network was constructed by thresholding this transition matrix, and analyzed using graph theory. The same approach was also applied to human rsfMRI data to examine the translational value of the findings in rats. A schematic illustration of these procedures is shown in Fig. 1.

**Figure 1.**
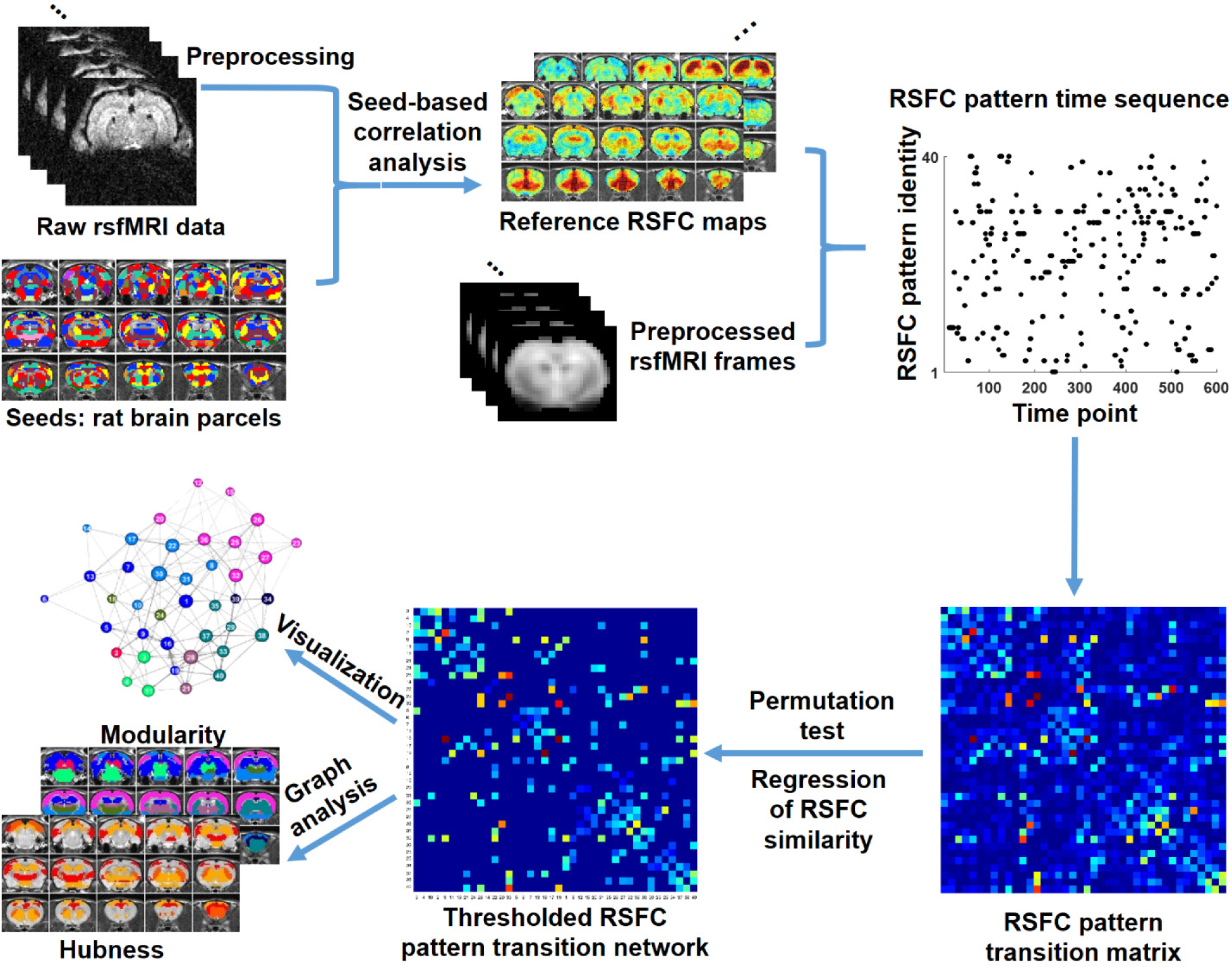
Schematic illustration of the data analysis pipeline.

### Characteristic RSFC patterns in the awake rat brain

An example of characteristic RSFC pattern was shown in Fig. 2, and the other 39 characteristic RSFC patterns were shown in Fig. 2-figure supplements 1. As the whole-brain parcellation scheme we adopted maximized within-parcel and minimized cross-parcel RSFC profile similarity, these 40 group-level seed-based RSFC maps represented a set of characteristic RSFC patterns in the awake rat brain, and were as the reference patterns. Notably, the number 40 was arbitrarily selected as an example of low-dimensionality parcellation of the rat brain. Similar analysis can be applied using other parcel numbers.

**Figure 2.**
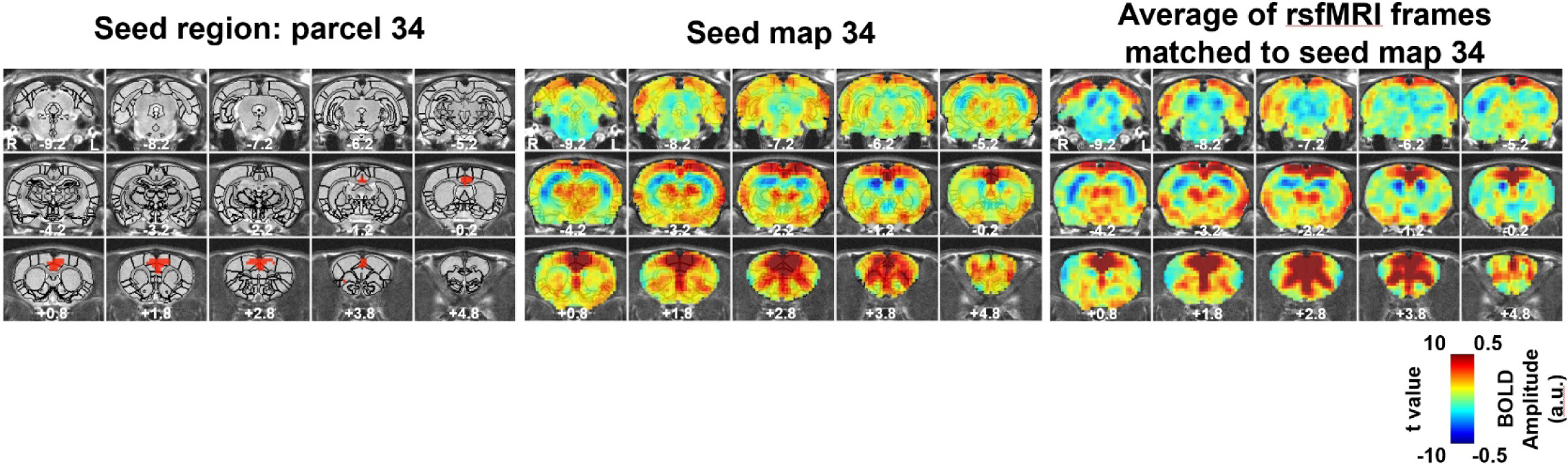
An example of RSFC spatial pattern. Left: seed region; Mid: RSFC pattern of the seed. Color bar indicates t values; Right: average of rsfMRI frames matched to the RSFC pattern. Color bar shows BOLD amplitude. Distance to bregma is listed at the bottom of each slice.

Fig. 2 (right panel) also showed the pattern of averaged rsfMRI frames that were matched to the reference RSFC pattern, which demonstrated high reminiscence between the BOLD co-activation pattern of single rsfMRI frames and the RSFC pattern it corresponded to.

### Reproducible temporal transitions between RSFC patterns

We first demonstrated that temporal transitions between RSFC patterns were highly reproducible at the group level. We randomly split all rats into two subgroups and obtained the transition matrix for each subgroup. Both matrices exhibited high similarity (Fig. 3a), reflected by a significant correlation (r = 0.86, p ≈ 0) between the corresponding off-diagonal entries. To control for the possible bias that transitions between similar RSFC patterns may have a higher chance to occur in both subgroups, which can inflate the reproducibility, we regressed out the spatial similarities between reference RSFC patterns from both transition matrices. The reproducibility remained high after regression, with a significant correlation value of 0.77 (p ≈ 0, Fig. 3b). Taken together, these results suggest that transitions between RSFC patterns are not random but follow specific temporal sequences in awake rats, and these transition sequences are not dictated by the similarity between RSFC patterns.

**Figure 3.**
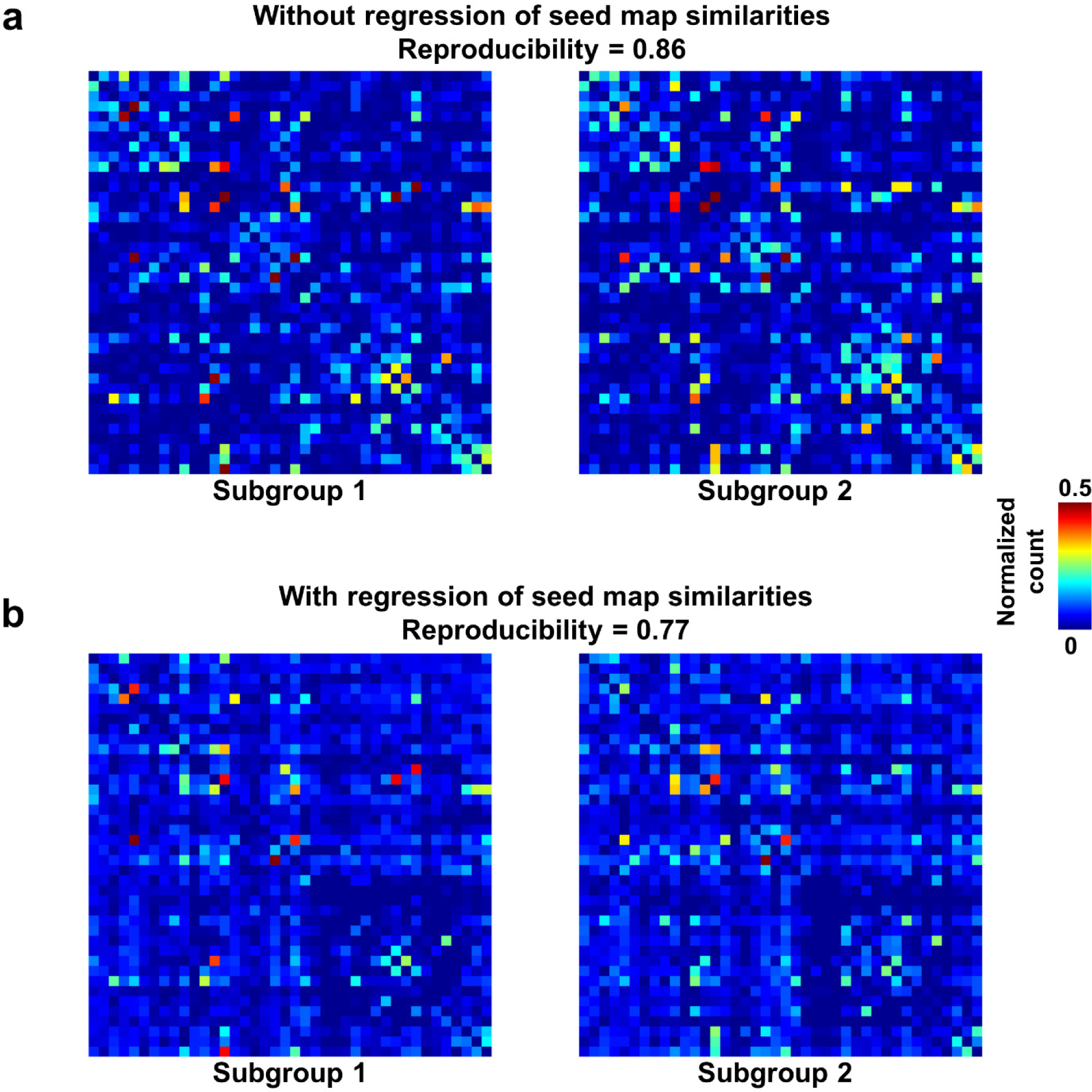
Reproducibility of the RSFC pattern transitions. a. RSFC pattern transition matrices of subgroup 1 and 2 without the regression of spatial similarities between reference RSFC patterns. b. RSFC pattern transition matrices of subgroup 1 and 2 with the regression of spatial similarities between reference RSFC patterns. Entries in each transition matrix were normalized to the range of [0, 1].

To further examine whether reproducible RSFC pattern transitions were dominated by a small portion of rats, we assessed the reproducibility of RSFC pattern transitions for each individual animal by computing Pearson correlation between each individual-level transition matrix and the group-level transition matrix. Fisher Z-transformed correlation values were then averaged across rats. Our data showed a significant individual-level reproducibility (mean (± SD) = 0.57 (± 0.14), p ≈ 0). These results collectively indicate that nonrandom RSFC pattern transitions are a characteristic feature in awake rats.

To rule out the possibility that RSFC pattern transitions were caused by head motion (Laumann TO et al. 2016), we conducted two additional analyses. First, we re-evaluated the reproducibility of RSFC transitions between two subgroups of rats with relatively high and low motion, respectively. Rats in the first subgroup all had the motion level below the median, quantified by FD. Rats in the second subgroup all had the motion level above the median. Transition matrices were obtained in these two subgroups, respectively. Comparing these two transition matrices yielded a reproducibility of 0.786 (with the regression of seed map similarities, Fig. 3-figure supplements 1), which is similar to the reproducibility assessed based on random grouping (0.77, Fig. 3b). In addition, the transition matrices from both subgroups were highly consistent with those in subgroups randomly divided (Fig. 3b). These results indicate that high reproducibility in RSFC pattern transitions was not due to head motion. In the second analysis, we directly compared the motion level between rsfMRI frames involved in RSFC pattern transitions versus those that were not in transitions. All rsfMRI frames analyzed were categorized into two groups. The first group included frames whose preceding and/or successive frame corresponded to a different RSFC pattern(s). The second group included frames whose preceding and successive frames were the same RSFC pattern. These two groups of rsfMRI frames showed consistent motion levels, quantified by their FD values (p = 0.44, two-sample t-test), again indicating that RSFC pattern transitions were not triggered by head motion.

### Within- and between-brain system transitions

Fig. 4 showed the group-level transition matrix thresholded using a permutation test (p < 0.05, FDR corrected). The order of rows/columns in the transition matrix was arranged based on the brain system (defined by different textures) that the seed region of each reference RSFC pattern belonged to. Transitions between RSFC patterns tended to occur within the same brain system, as shown by a relatively denser distribution of near-diagonal nonzero elements in the matrix. In addition to prominent within-system RSFC pattern transitions in most brain systems, cross-system transitions such as striatal-thalamic, striatal-somatosensory, striatal-prefrontal, striatal-hippocampal, hippocampal-amygdala, amygdala-motor transitions were observed.

**Figure 4.**
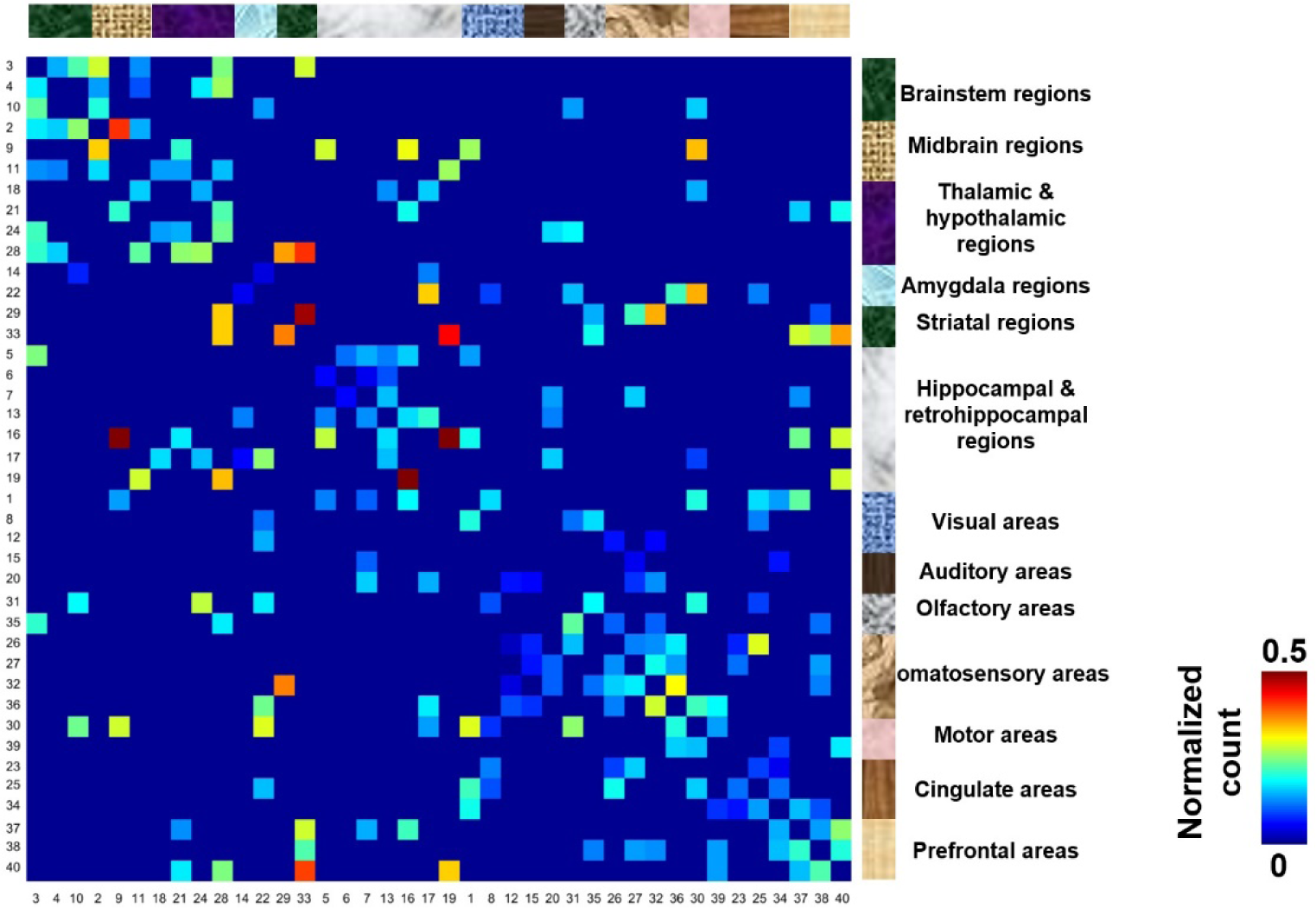
Thresholded group-level RSFC pattern transition matrix after the regression of RSFC pattern similarities. Rows/columns are arranged based on the brain system (defined by different textures) the seed regions belonged to. Numbers next to/below rows/columns correspond to the seed map number in Fig. 2 and Fig. 2-supplements 1.

### Organization of RSFC pattern transitions

A directed weighted graph of the RSFC pattern transition network was constructed based on the group-level thresholded transition matrix (Fig. 4), as shown in Fig. 5. The number of edges was 242, yielding a connection density of 15.5%. The transition network exhibited a prominent community structure with nine modules identified using the Louvain community detection algorithm (Vincent DB *et al.* 2008), suggesting that RSFC patterns belonging to the same modules had a higher probability to transit between each other than RSFC patterns across modules. The corresponding seed regions of RSFC patterns were color coded based on the community affiliations (Fig. 5 inlet). Module 1 primarily covered hippocampal and retrohippocampal networks as well as caudal visual networks. Module 2 included caudal midbrain networks. Module 3 was comprised of brainstem and rostral midbrain networks. Module 4 covered rostral visual, amygdala, hypothalamic as well as motor and olfactory networks. Module 5 was dominated by auditory and somatosensory networks. Module 6 captured posterior ventral thalamic networks. Module 7 included anterior thalamic networks. Module 8 covered striatal and prefrontal networks. Module 9 mainly included anterior cingulate cortex network. Networks from the same system usually fell into the same community, again indicating transitions between RSFC patterns frequently occurred within the same brain system. However, networks from different systems were also observed in the same modules, which highlights the importance of cross-system transitions.

**Figure 5.**
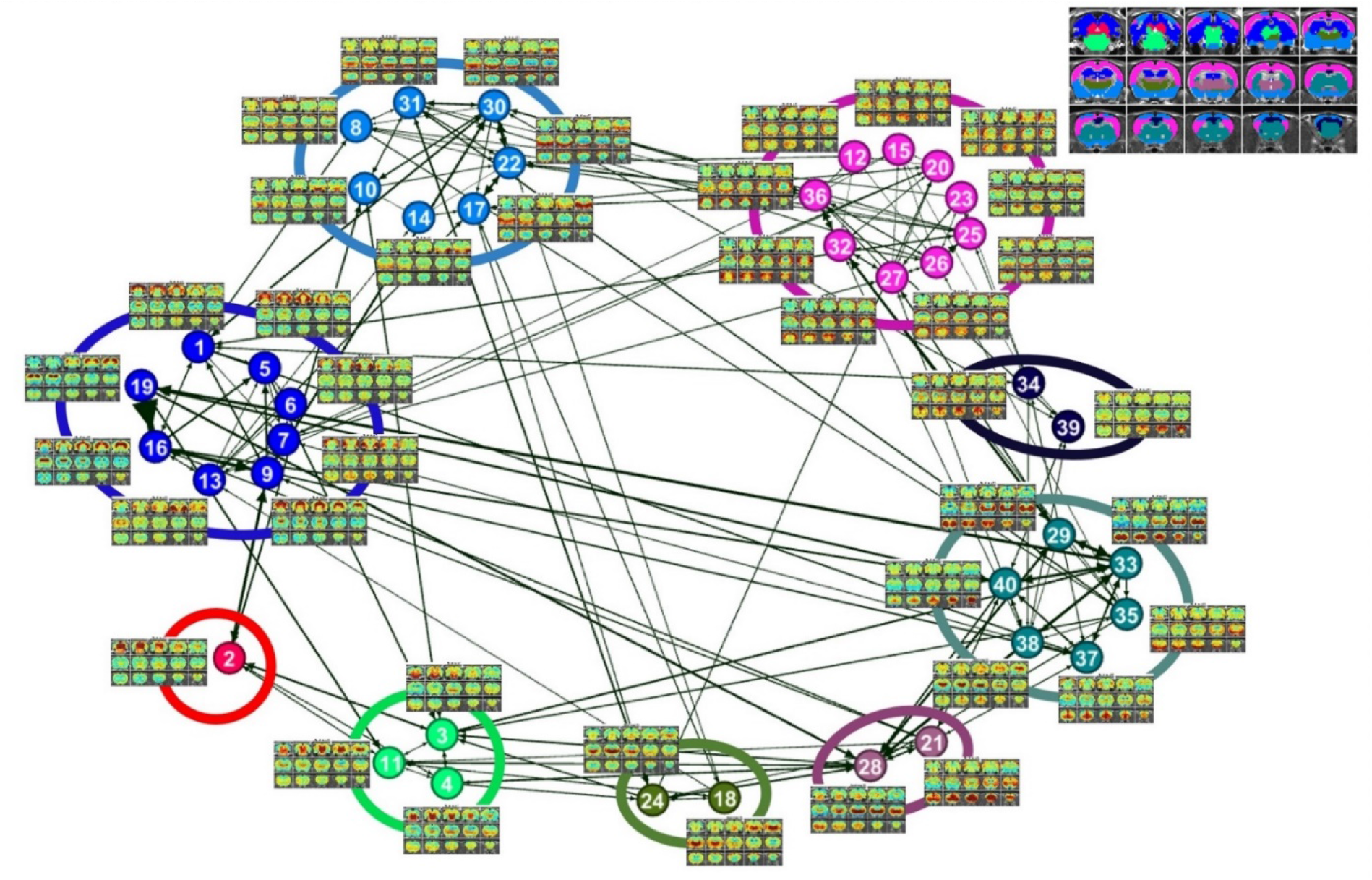
Community structures of the RSFC pattern transition network. The thresholded transition matrix in Fig. 4 was used as the adjacency matrix to generate a directed weighted graph. The layout of nodes was based on a force-field algorithm (Jacomy M et al. 2014). The node number corresponds to the seed map number in Fig. 2 and Fig. 2-supplements 1. Inlet: seed regions of all RSFC patterns colored coded based on the community affiliations of nodes (i.e. RSFC patterns).

By quantifying the node-specific graph measures of node strength, betweenness centrality, characteristic path length and local clustering coefficient, hub nodes (i.e. pivotal RSFC patterns) in the graph were identified. Six RSFC patterns were identified as hubs (hub score ≥ 3. Fig. 6a) including the networks of retrosplenial cortex, dorsal superior and inferior colliculi, hippocampus, anterior ventral thalamus, striatum and motor cortex. Fig. 6b also showed the seed regions of RSFC patterns with hub score ≥ 1, color coded based on the hub score.

**Figure 6.**
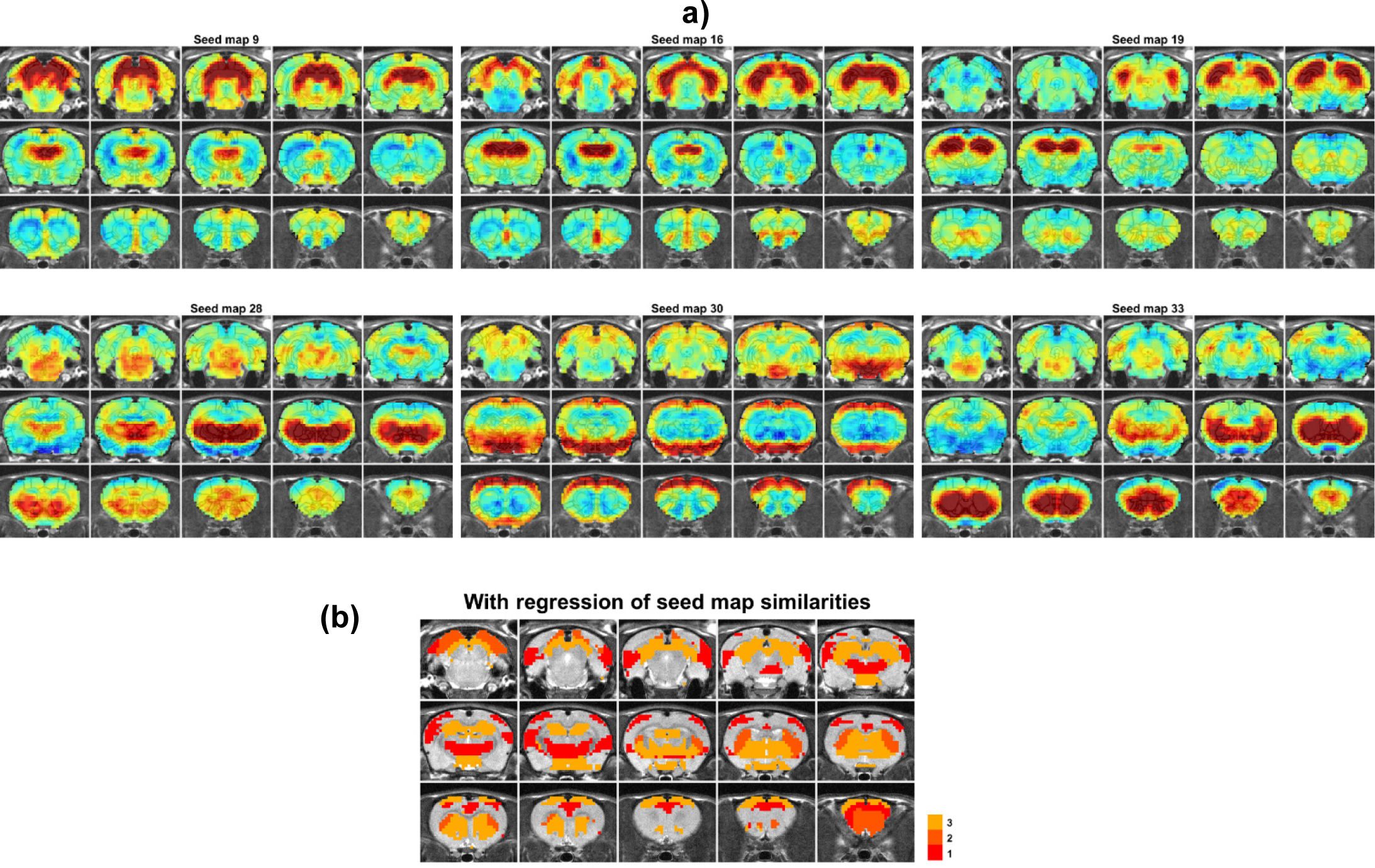
a) Hub patterns in RSFC transitions. b) Seed regions of RSFC patterns with hub score ≥ 1, color coded based on the hub score.

Fig. 7 showed RSFC pattern transitions of four representative hubs, demonstrating the pivotal role of these patterns in RSFC temporal transitions. The majority of RSFC pattern transitions between hubs and other RSFC patterns were bidirectional. Fig. 7a showed transitions of the hub network of the superior and inferior colliculi with the networks of the periaqueductal gray, hippocampus, dorsal thalamus, hypothalamus, caudal visual cortex and motor areas. Fig. 7b demonstrated the transitions between the hippocampus network (hub) and the RSFC networks of the superior and inferior colliculi, hippocampus/retrohippocampus, caudal visual cortex, as well as prefrontal and orbital cortices. Fig. 7c displayed the transitions between the hub network of the anterior ventral thalamus and the RSFC networks of the brainstem, midbrain, dorsal CA1, dorsal thalamus, posterior ventral thalamus, dorsal caudate-putamen (CPu), olfactory tubercle, and orbital cortex. Fig. 7d illustrated the transitions between the hub network of the ventral CPu, and the RSFC networks of the brainstem and olfactory tubercle, as well as infralimbic, prelimbic, and orbital cortices. Taken together, these results indicate that hub RSFC patterns were centralized patterns that play a pivotal role in transitions with other RSFC patterns involving multiple brain systems.

**Figure 7.**
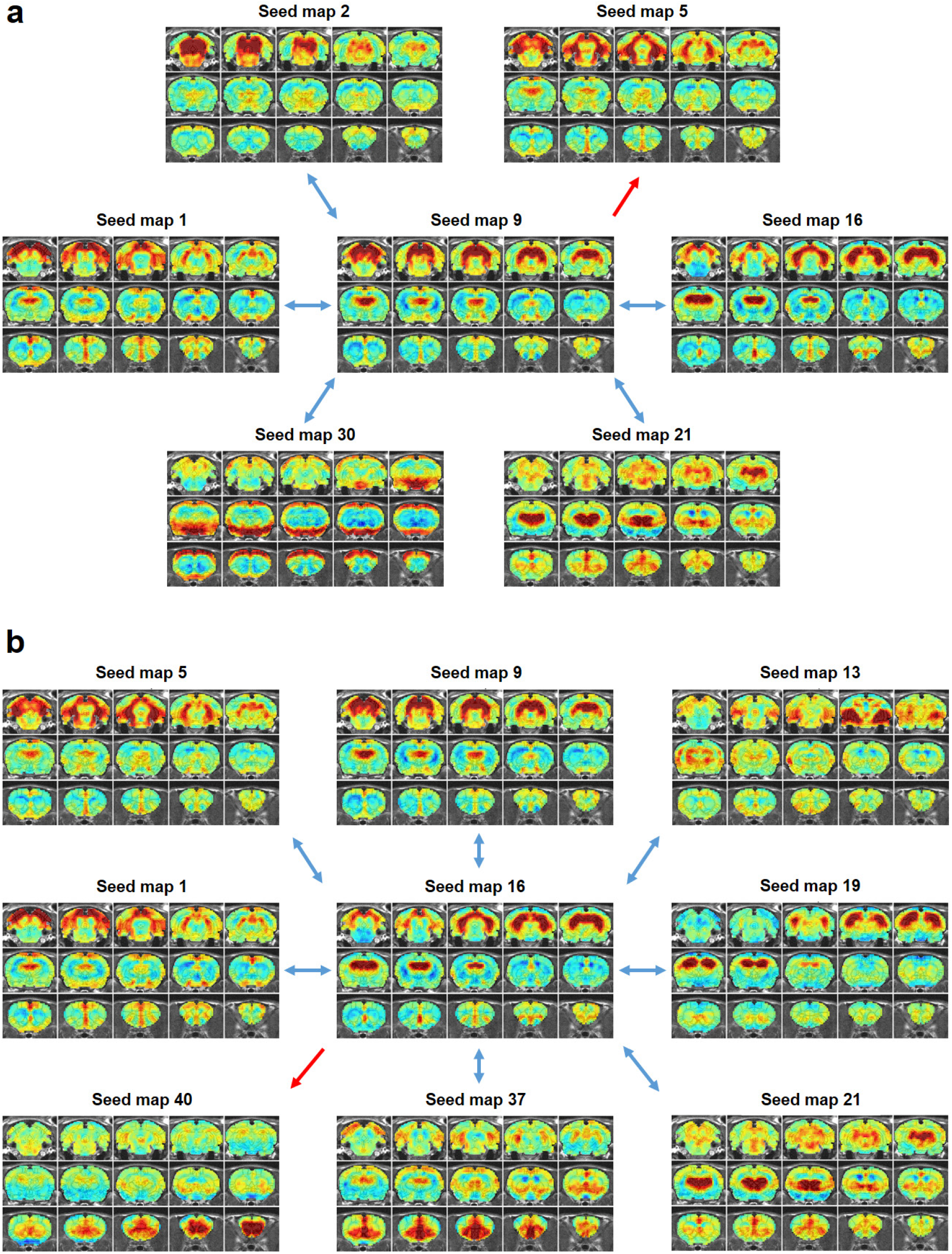
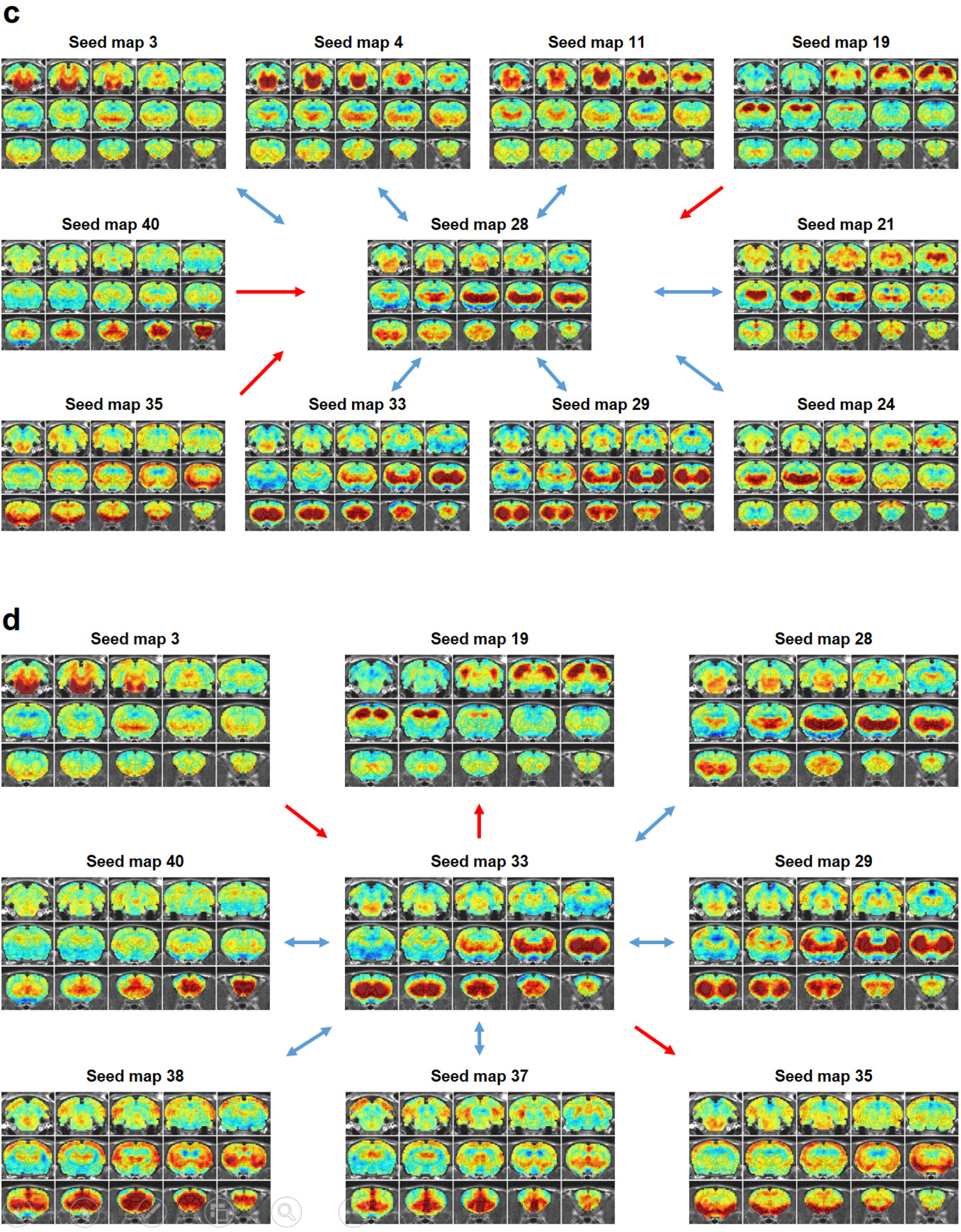
Transition patterns of hub networks. Blue arrows denote bidirectional transitions between RSFC patterns. Red arrows denote unidirectional transitions between RSFC patterns.

### RSFC pattern transitions in humans

To assess whether temporal transitions between RSFC patterns are also nonrandom in humans, we applied the same analysis to rsfMRI data from 812 human subjects in the HCP. Each frame was matched to one of 333 characteristic RSFC patterns defined by a well-established RSFC-based parcellation in humans (Gordon EM *et al.* 2016), and the number of transitions between every two RSFC patterns was counted for each subject. All subjects were then randomly split into two subgroups (406 subjects in each subgroup). The reproducibility between two subgroups was 0.9955 (without regression of seed map similarities, Fig. 8a), and 0.9954 (with the regression of seed map similarities, Fig. 8b). To assess the reproducibility at the individual level, the correlation between the transition matrix of each individual subject versus the group-level transition matrix was calculated. The mean correlation (±SD) across all subjects was 0.60 (±0.05). All these results were highly consistent with our findings in awake rats, suggesting that nonrandom transitions between RSFC patterns are conserved across species and might represent a characteristic feature of the mammalian brain.

**Figure 8.**
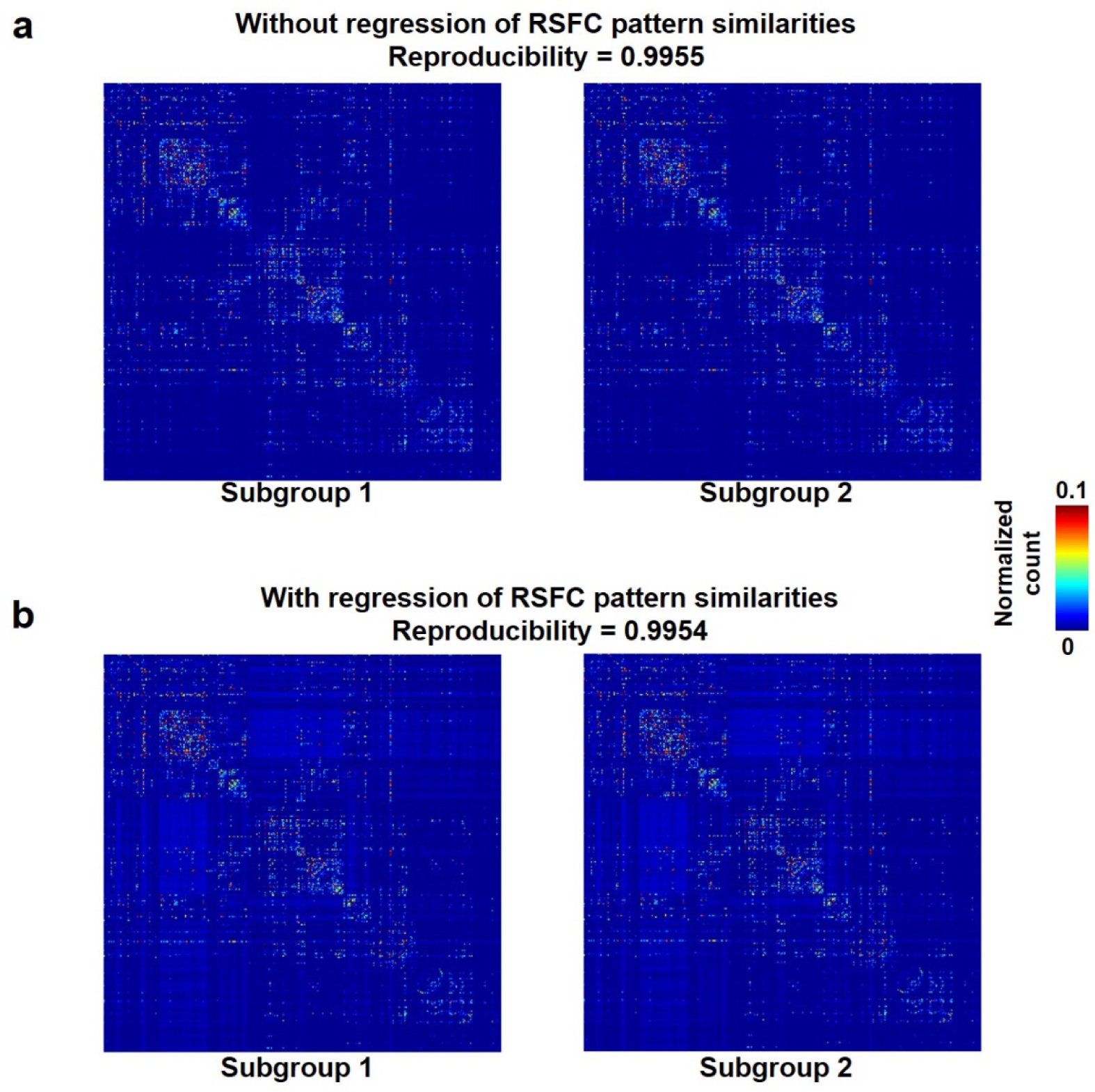
Reproducibility of temporal transitions between 333 characteristic RSFC patterns in humans. The transition matrices of two randomly divided subgroups (a) without regression of RSFC pattern similarities (b) with the regression of RSFC pattern similarities.

## Discussion

In the present study, we investigated temporal sequential transitions between intrinsic brain activity patterns in the awake rat brain. We showed that transitions between RSFC patterns exhibited high reproducibility across animals and were significantly above chance (Figs. 3 and 4). In addition, the RSFC pattern transition network was constructed (Fig. 5a) using the thresholded group-level transition matrix (Fig. 4), and its topological organization including the community structure (Fig. 5) and hubness (Fig. 6) was evaluated. Moreover, the transitions of representative hub RSFC patterns were demonstrated (Fig. 7). Importantly, non-random RSFC pattern transitions were also observed in humans (Fig. 8), highlighting the translational value of this feature. Taken together, the present study for the first time characterized the temporal sequence between successive brain connectivity configurations. It demonstrates that characteristic RSFC patterns do not fluctuate in a random manner, but follow specific orders. These results revealed the temporal relationship between characteristic RSFC spatial patterns in both rats and humans, and has provided new insight into understanding the spatiotemporal dynamics of spontaneous activity in the mammalian brain.

### Method to unveil the temporal relationship between characteristic RSFC patterns

Although it has been well recognized that RSFC is dynamic in nature (Hutchison RM, T Womelsdorf, EA Allen*, et al.* 2013), previous studies in this research line generally focused on revealing the spatial features of recurring RSFC patterns. Meanwhile, we have relatively sparse knowledge in the temporal relationship between characteristic RSFC patterns at the resting state (Majeed W *et al.* 2011; Zalesky A *et al.* 2014; Vidaurre D et al. 2017). To bridge this gap, here we systematically investigated temporal transitions between RSFC patterns.

To address this issue, we first need a set of representative RSFC patterns in the awake rat brain. Since the rat brain has ~6000 brain voxels at our spatial resolution (0.5×0.5×1 mm^3^), in principle we can have in total ~6000 RSFC profiles. However, elucidating temporal sequences between such a large number of RSFC patterns is not only computationally intensive, but also not necessary as many of these patterns are highly similar to each other. To obtain a survey of characteristic RSFC patterns, we adopted a RSFC-based parcellation of the awake rat brain (Ma Z *et al.* 2016). In this scheme, all ~6000 voxels were clustered into 40 parcels based on the similarity of their RSFC patterns, so that brain voxels’ RSFC profiles were similar within each parcel but dissimilar between parcels (Ma Z *et al.* 2016). Notably, these parcels were highly reproducible between animals and exhibited high with-parcel homogeneity (Ma Z *et al.* 2016). Therefore, RSFC patterns obtained based on these parcels provided a representation of all ~6000 RSFC patterns.

To examine the temporal relationship between these characteristic RSFC patterns, we adapted a recently developed method showing that BOLD co-activation patterns of rsfMRI frames well correspond to their instantaneous RSFC patterns (Liu X *et al.* 2013; Liu X and JH Duyn 2013). This notion has been demonstrated in both humans, as well as in awake and anesthetized rats (Liang Z *et al.* 2015). Using this notion, each rsfMRI frame was corresponded to one of the 40 characteristic RSFC patterns based on the spatial similarity to the frame’s BOLD co-activation pattern. This step resulted in a time sequence of RSFC patterns, which allowed us to systematically investigate the temporal transitions between these RSFC patterns.

### Nonrandom temporal transitions between RSFC patterns

Our data showed that temporal transitions between RSFC patterns were highly reproducible, reflected by significant reproducibility between randomly divided subgroups. In addition, these reproducible transitions were not dominated by a small portion of animals, evidenced by highly significant reproducibility at the individual level. It should be noted that the reproducibility assessment using the split-group method relies on the assumption that RSFC pattern transitions were independent. However, transitions between spatially more similar RSFC patterns may occur at a higher chance in both subgroups. This potential systematic bias can artificially increase the reproducibility between the two subgroups. To control for this factor, similarities between RSFC patterns were regressed out in the transition matrices of both subgroups, and we found that the reproducibility of transitions remained high. These data show that transitions between RSFC patterns were robust. In addition, using permutation tests, we identified a number of transitions between RSFC patterns that were statistically above chance. We also ruled out the possibility that RSFC pattern transitions were driven by head motion. Consistent transition matrices were obtained in two subgroups of animals with low (below median) and high (above median) motion levels. In addition, no difference in head motion was observed between rsfMRI frames involved and not involved in transitions. Taken together, these results provide strong evidence indicating that RSFC patterns do not transit from/to each other in a random manner, but follow specific temporal sequences. These results well agree with a report that spontaneous activity from ensembles of simultaneously recorded neurons is characterized by ongoing spatiotemporal activity patterns (Mazzucato L et al. 2015), which recur during all trials, and transitions between patterns can be reliably extracted using a hidden Markov model (Mazzucato L *et al.* 2015).

### RSFC pattern transitions in humans

To examine whether nonrandom RSFC pattern transitions we observed were only a feature in the awake rat brain, we investigated RSFC pattern transitions in humans by applying the same analysis approach to rsfMRI data from the HCP. We found that, like rats, transitions between RSFC patterns were also nonrandom in humans, evidenced by highly consistent transition matrices between two randomly divided subject subgroups. This result well agrees with a recent study showing that dynamic switching between human brain networks is not random (Vidaurre D *et al.* 2017). Interestingly, the split-group reproducibility rates were somewhat higher in humans than those observed in rats (both with and without regression of RSFC pattern similarities). This difference is likely due to much more human data used (406/406 human subjects v.s. 20/21 rats in each subgroup), which would average out larger amount of individual variability. This concept can be further supported by comparable reproducibility rates if we randomly picked 20 human subjects for each subgroup (reproducibility = 0.91 for human data v.s. reproducibility = 0.86 for rat data), as well as similar reproducibility at the individual level (human data: 0.6(± 0.05); rat data: 0.57(±0.14)). Collectively, these results suggest that nonrandom transitions between characteristic RSFC patterns are not merely a feature in rodents, but conserved across species in both humans and rats. Such findings highlight the translational utility of the analysis applied in the present study, which might shed light onto comparative neuroanatomy. Our results have also provided new insight into understanding the spatiotemporal dynamics of spontaneous activity in the mammalian brain.

### Transitions between RSFC patterns within and across brain systems

We found that transitions between RSFC patterns occurred frequently between networks from the same brain system (Fig. 4). This result might be attributed to the factor that seed regions of networks in the same brain system typically subserve similar brain function. In addition, regions in the same brain system are usually strongly connected with each other (Liang Z et al. 2013), and thus transitions between their RSFC patterns can frequently occur.

Our data also showed prominent cross-system transitions (Fig. 4). For instance, switching between striatal networks and somatosensory/prefrontal cortical networks frequently occurred. Such cortical-subcortical system transitions might rely on the structural basis of corticostriatal projections identified in the rat brain (Paxinos G 2015). We speculate that bidirectional transitions between striatal and somatosensory/prefrontal RSFC networks might indicate the presence of both “bottom-up” and “top-down” processing involving high-order cortical and low-order subcortical regions at rest (Gurney KN et al. 2015; Piray P et al. 2016). In addition, significant transitions from striatal to thalamic/hippocampal RSFC networks indicate a close relationship between these subcortical systems, which can be further supported by strong RSFC between the CPu and thalamus found in the awake rat brain (Liang Z *et al.* 2013). Taken together, these results show non-trivial transitions between RSFC patterns within and across systems in the awake rat brain, and such transitions might play a critical role in coordinating spontaneous brain activity in separating brain systems.

### Organization of the RSFC pattern transition network

A graph characterizing the transition network between RSFC patterns was constructed with each node representing a characteristic RSFC pattern and each edge denoting a statistically significant transition relationship between two nodes. We investigated the topological organization of this weighted directed graph including its community structure and hubness. The transition network exhibited a prominent community structure evidenced by a high modularity, indicating that the global transition network between RSFC patterns was organized in a non-trivial manner.

We also identified several hub RSFC patterns (Fig. 6) and scrutinized their transitions with other RSFC patterns (Fig. 7). Hub patterns are central nodes in the RSFC transition network which play a pivotal role in transitions from/to other RSFC patterns. We found that the hippocampus RSFC network was pivotal to the transitions to the superior and inferior colliculi networks, as well as visual, prefrontal and orbital cortical networks. A recent study has shown that low-frequency hippocampal–cortical activity drives brain-wide rsfMRI connectivity, highlighting the pivotal role of the hippocampus in RSFC transitions (Chan RW et al. 2017). In addition, Xiao and colleagues demonstrate that hippocampal spikes are associated with calcium cortical co-activation patterns in the visual and cingulate cortical regions (Xiao D et al. 2017), consistent with our observation of transitions between hippocampal networks and visual/cingulate networks (Fig. 7). Interestingly, it has also been reported that the hippocampus interacts with multiple cortical and subcortical regions in the form of sharp wave ripples (Logothetis NK et al. 2012), and the onset of such ripples was found to be controlled by the propagating signals from the cortex to hippocampus (Molle M et al. 2006; Hahn TT et al. 2012; Roumis DK and LM Frank 2015). These results agree with the bidirectional transitions between hippocampal and cortical networks found in the present study (Fig. 7). Although sharp wave ripples in the hippocampus and RSFC pattern transitions might be from different signal sources, the centralized role of the hippocampus is shared by these two forms of hippocampal-cortical information flow.

In accordance with our previous report that anterior ventral thalamus is a critical hub in the rat functional brain network (Ma Z *et al.* 2016), the RSFC pattern of anterior ventral thalamus was also a hub in the network of RSFC pattern transitions. A recent study investigating the relationship between single neuron spiking activity and brain-wide cortical calcium dynamics found that thalamic spikes could both predict and report (i.e. firing before and after) various types of large-scale cortical activity patterns, which are supported by slow calcium activities (<1 Hz) (Xiao D *et al.* 2017). These data indicate a pivotal role of the thalamus in transitions between distinct spontaneous brain activity patterns. This result can be further supported by the finding that the ventral thalamus can recruit long-range cortical and subcortical networks and initiate their interactions through low-frequency (1 Hz) activity (Leong AT et al. 2016), and the thalamus can facilitate diverse and brain-wide functional neural integration in a specific spatiotemporal manner (Liu J et al. 2015; Leong AT *et al.* 2016).

Further, our data revealed a hub of the RSFC pattern of the ventral CPu. As a part of the striatum, CPu is linked to multiple corticostriatal projections (Paxinos G 2015), and it might play a centralized role in transitions involving multiple cortical RSFC patterns (Lee K et al. 2017). Taken together, these data indicate that hub RSFC patterns might play a centralized role linking multiple brain systems and might be critical for us to understand how activities from different brain systems are integrated to maintain normal brain function.

### Potential limitation

One limitation of the present study is that single rsfMRI frames could exhibit features of more than one RSFC pattern. It should be noted that corresponding a rsfMRI frame to its most similar reference RSFC pattern is only an approximation for the purpose of investigating spatiotemporal dynamics of spontaneous brain activity. To mitigate this issue, we set a minimal threshold (correlation coefficient > 0.1, p < 10^−13^) to remove rsfMRI frames that were not similar to any of 40 reference RSFC patterns (e.g. rsfMRI frames dominated by noise), and ensured that the similarity between each rsfMRI frame and the RSFC pattern it corresponded to was statistically significant after Bonferroni correction (p < 0.05/40834 rsfMRI volumes ≈ 10^−6^). 89.9% of total rsfMRI volumes met this criterion, indicating that reference RSFC patterns indeed captured most spontaneous brain activity patterns in the awake rat brain.

## Conclusions

In conclusion, the present study investigated temporal transitions between spontaneous brain activity patterns in the awake rat and human brain. The temporal sequence of framewise RSFC patterns was obtained, and reproducible transitions between RSFC patterns in the sequence were identified. Using graph theory analysis, our study further revealed community structures and hubness of the RSFC pattern transition network. This study has opened a new avenue to investigating the spatiotemporal relationship of spontaneous activity in the mammalian brain.

## Acknowledgments

The present study was partially supported by National Institute of Neurological Disorders and Stroke Grant R01NS085200 (PI: Nanyin Zhang, PhD) and National Institute of Mental Health Grant R01MH098003 and RF1MH114224 (PI: Nanyin Zhang, PhD). Part of this research was conducted using the high-performance computing resources provided by the Institute for CyberScience at the Pennsylvania State University (https://ics.psu.edu). Human data were [in part] provided by the HCP, WU-Minn Consortium (Principal Investigators: David Van Essen and Kamil Ugurbil; 1U54MH091657) funded by 16 NIH Institutes and Centers that support the NIH Blueprint for Neuroscience Research; and by the McDonnell Center for Systems Neuroscience at Washington University.

